# Insight into human photoreceptor function: modeling optoretinographic responses to diverse stimuli

**DOI:** 10.1101/2025.02.28.639986

**Authors:** Denise Valente, Kari V. Vienola, Robert J. Zawadzki, Ravi S. Jonnal

## Abstract

Optoretinography is an emerging method for detecting and measuring functional responses from neurons in the living human retina. Its potential applications are significant and broad, spanning clinical assessment of retinal disease, investigation of fundamental scientific questions, and rapid evaluation of experimental therapeutics for blinding retinal diseases. Progress in all these domains hinges on the development of robust methods for quantifying observed responses in relation to visible stimuli. In this work, we describe a novel optoretinographic imaging platform–full-field swept-source optical coherence tomography with adaptive optics, measure cone responses in two healthy volunteers to a variety of stimulus patterns, and propose a simple model for predicting and quantifying responses to those stimuli.

**Teaser:** The function of human retinal neurons is quantified by the light they reflect, opening a new frontier in visual health

## 1 Introduction

Photoreceptors are specialized neurons responsible for phototransduction, a biochemical process that initiates vision. Therefore, the assessment of these cells’ function is a critical step when evaluating vision. Blinding diseases whose pathogenesis primarily involves photoreceptors include age-related macular degeneration (AMD), retinitis pigmentosa (RP), and other inherited retinal degenerations (IRDs) such as achromatopsia, pattern dystrophy, and Stargardt’s disease. AMD is the most prevalent of these, affecting nearly 200 million people worldwide, with projections estimating that this number will reach 300 million within the next two decades (*1*), making it the leading cause of irreversible blindness globally. RP and other IRDs affect an additional 5 million individuals (*2, 3*).

At present, the predominant means of assessing retinal function are subjective–tests such as visual fields, visual acuity (VA), and contrast sensitivity. Less frequently employed are objective measures like the electroretinograms (ERG) and multifocal (mf)ERG. Despite their clinical utility, all of these methods have inherent limitations. VA tests only assess central function, and all subjective tests are affected by sources of noise such as fatigue, attention, and learning effects. While ERG testing is objective, it is slightly invasive and lacks spatial resolution. Conversely, mfERG offers spatial localization up to one degree but can be a demanding test, requiring stable fixation for ten minutes or more. While perimetry and mfERG provide some spatial localization, they do not provide accompanying information about the retinal structure in those locations.

Photoreceptor health can also be assessed using structural imaging. Optical coherence tomography (OCT) offers high-resolution three-dimensional imaging of the laminar structure of the living human retina. It has become a standard of ophthalmic care by providing structural biomarkers of retinal disease progression (*4*). However, although OCT-based structural endpoints are predictive of changes in visual function, they often only detect those changes after significant damage has been observed (*5*). Moreover, in the context of therapies designed to protect or restore visual function, the structural changes defining an endpoint may represent unrecoverable vision loss.

Structural OCT imaging relies on the amplitude of the OCT signal. However, since OCT is based on interferometry, it also provides access to the phase of the measured interference fringes. Motion of structures in the retina, even when much smaller than the OCT axial resolution or the wavelength of the imaging light, cause predictable changes in the OCT phase (*6*). Thus changes in the OCT phase can be used to infer motion relative to the retina, after compensation of movement of the eye and head using statistical (*7*) or direct (*8*) methods.

In recent years, combination of phase-sensitive imaging with visible stimuli has led to the discovery of nanometer-scale light-evoked changes in the photoreceptor outer segments consisting of reproducible patterns of contraction and elongation that mostly scale with the stimulus dose (*9–16*). This signal has emerged as a favored tool in the broader set of methods used to measure retinal neural function optically, which has been called optoretinography (ORG).

Besides phototransduction, which permits the generation of a visual signal, another functional process critical for photorecetpors is the visual cycle, important for their homeostasis and photopigment replenishment. Both phototransduction and the visual cycle are complex, consisting of multiple enzymatic and energetic steps. Disease-related disruption of the different steps may result in changes to different aspects or features of the normal ORG response. Therefore, it may be useful to devise methods for the quantitative description of these features.

In this paper we describe a novel optoretinographic imaging platform consisting of a fullfield (FF), swept-source (SS), optical coherence tomography (OCT) system with adaptive optics (AO) and a visual stimulation channel. We used the system to measure ORG responses from two healthy subjects under a variety of stimulus conditions. We propose a four-parameter model that appears to describe the responses of cones to single-flash stimuli. We then discuss the extent to which the model can be used to analyse responses to more complex stimuli, such as multiple flashes and flashes delivered against adapting backgrounds, and the model’s implications for the sensitivity and bandwidth of ORG measurement.

## 2 Results

Clear photoreceptor responses were observed in both subjects in each stimulus condition: single flashes, paired flashes, pulse trains, and adapting backgrounds. Fits to the primary model

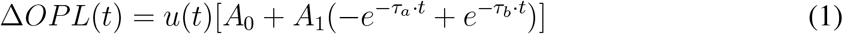

and its variants (described in the Materials and Methods section below) were qualitatively good and had low residual error. Results of the various stimulus conditions are described below.

### 2.1 Single flash

A representative measurement (subject 1, 8% bleach) is shown in Fig. 1 (a), along with the model fit. Optimal fit of the model was achieved with *A*_0_ = −36.4 nm, *A*_1_ = 200.8 nm, *τ*_*a*_ = 9.5 s^−1^, and *τ*_*b*_ = 0.07 s^−1^. Fits to all measurements were similarly qualitatively good, with *R*^2^ > 0.95 and root mean square (RMS) error < 22 nm for all trials (see Supplementary Table S1). Fig. 1 (b) illustrates the good fit during the initial OS contraction and start of elongation. The four parameters of the model have distinct effects on the shape of the curve. These effects are illustrated in Fig. 1 (c) for *A*_0_ and Fig. 1 (d) for *A*_1_, *τ*_*a*_, and *τ*_*b*_.

**Figure 1.**
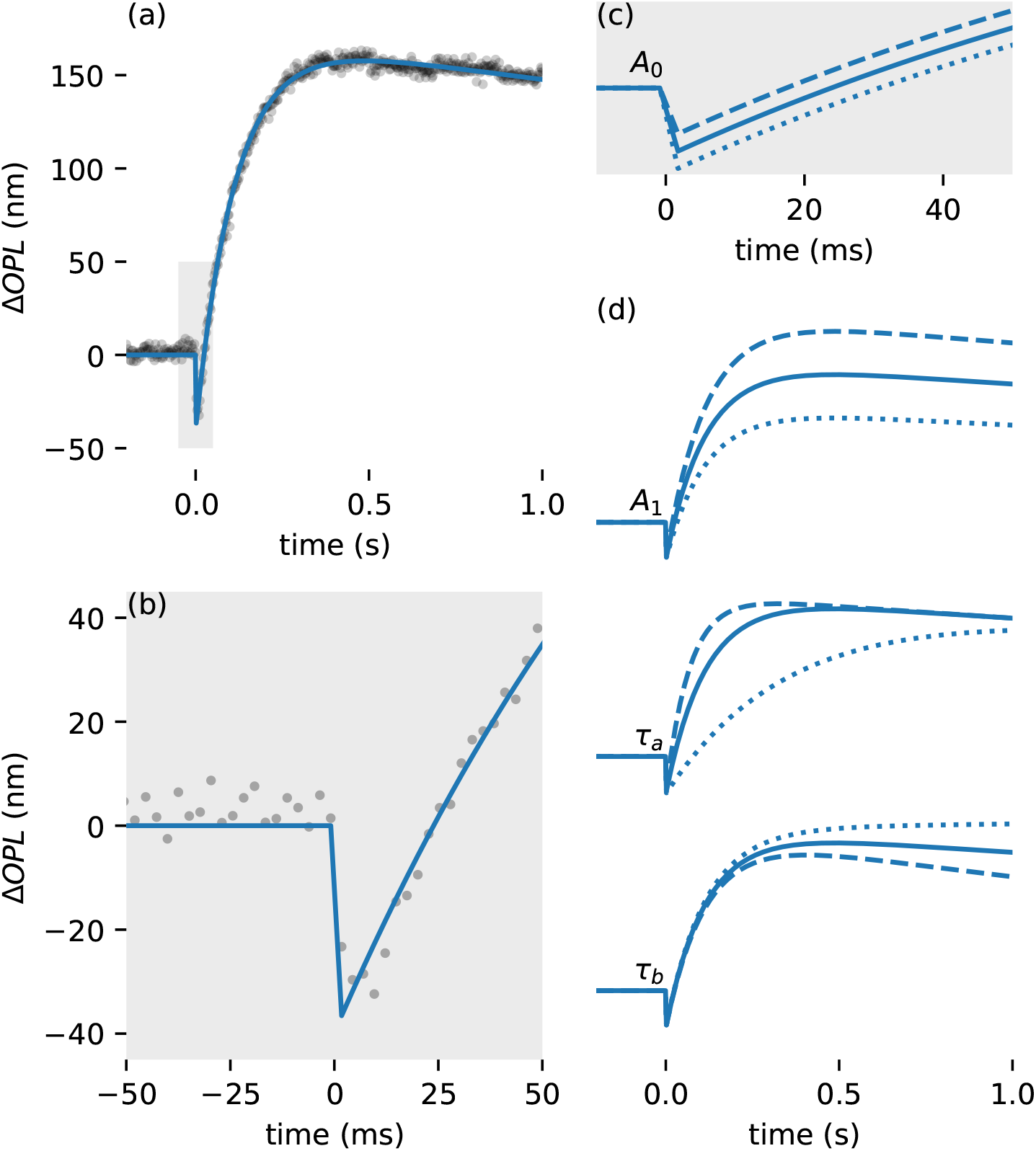
Representative measurement and fit, and the effects of model parameters on response. (a) Representative measurement (subject 1, 8% bleach) plotted (gray circles), along with model fit (blue line). The gray box indicates the portion of the fitted response shown in (b). (b) Early portion of response and fit shown in detail. (c) Illustration of the effect of parameter *A*_0_ on the ORG response, mainly seen in the amplitude of the initial OS contraction. (d) Illustrations of the effects of parameters *A*_1_, *τ*_*a*_, and *τ*_*b*_ on the ORG response. Dashed lines indicate more positive values, while dotted lines indicate less positive ones.

In single flash trials, measurements were performed in two subjects with various bleaching levels (Fig. 2) and fitted to the proposed model (Eq. 1). Measurements of the first subject were collected over 3 s after stimulus, while the second was measured for just 1 s. The reduced acquisition time in the second subject was motivated by the subsequent reductions in data transfer and processing times and data storage requirements. As shown below, this reduced time window appears primarily to impact estimation of the recovery rate parameter (*τ*_*B*_).

**Figure 2.**
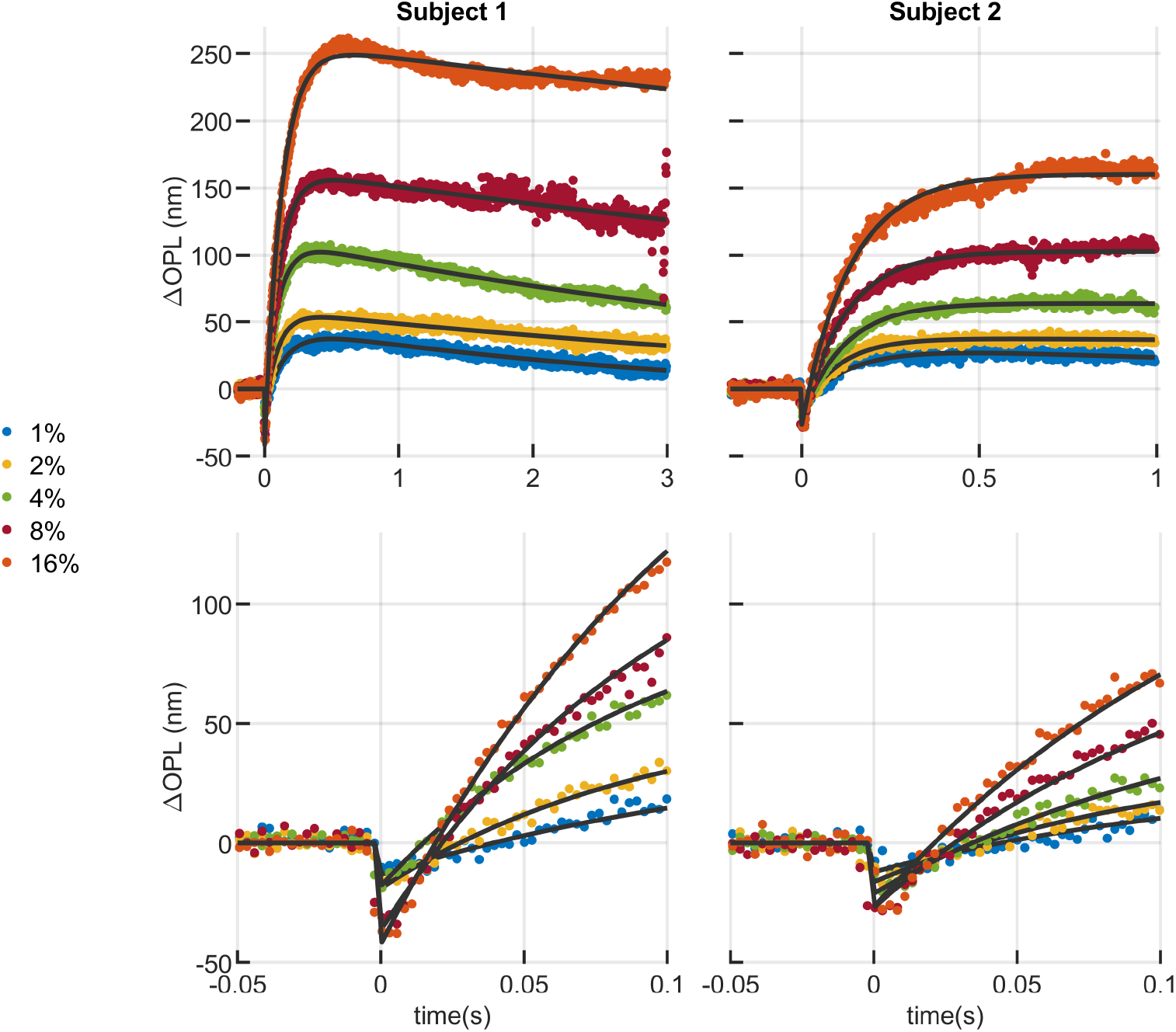
Curve fitting of ORGs of two subjects in response to a single flash of 10 ms and different bleaching levels. (top) The model provided visually good fits over a wide range of bleaching levels, whether ORG responses were measured over 3 s (subject 1, left) or 1 s (subject 2, right). In both subjects, some aspects of the response were observed to vary monotonically with dose, such as the maximum excursion (Δ*OPL*_*max*_) and slope of elongation between the flash and first ~100 ms increased with increasing bleaching fraction. Considerable variation between subjects was observed for given doses. This is most visible in the 16% trials, where the maximum OS excursion of subject 1 is at least 50% higher than that of subject 2. (bottom) The model provided qualitatively good fits to the early contractile portion of the response as well, though similar inter-subject variability is visible. The amplitude of the OS contraction appears to scale with dose, as well as the initial slope of elongation. The model has a discontinuity at *t* = 0 and thus does not capture features of the negative-going portion of the curve in the first 5 to 8 ms. This portion represents only 2 to 3 datapoints at the 400 Hz volume rate. This rate provides insufficient temporal resolution to model the initial contraction.

The effect of sampling duration on parameter estimation is illustrated in Supplementary Fig. S1, where the resulting percentage error for each parameter is depicted as a function of the observation window. An arbitrary level of 10% is plotted as a horizontal line in each of the four plots. It can be observed that *A*_0_ exhibits small variability, even with very short observation windows. However, the other parameters demonstrate a need for longer observation times. It is apparent that *A*_1_,*τ*_*A*_ and *τ*_*B*_ require at least 1.05 s, 1.85 s, and 2.85 s, respectively, within this bleaching range, although *A*_1_ and *τ*_*A*_ approach the 10% error level earlier for most bleaching levels. The most significant fluctuations are observed in *τ*_*B*_, associated with the rate at which the OS returns to its baseline length. The apparent convergence of error toward zero in the case of *τ*_*B*_ is an artifact of the analytical method, since error is defined as a proportional difference from the 3 s estimate.

The fitted parameters resulting from both subjects’ data using Eq. 1 are presented in Fig. 3. Our initial observation was that *A*_0_ exhibited linear behavior on a semi-logarithmic scale, decreasing (i.e., showing a higher negative amplitude) with dose. Thus, for *A*_0_, a log-linear model was investigated:

**Figure 3.**
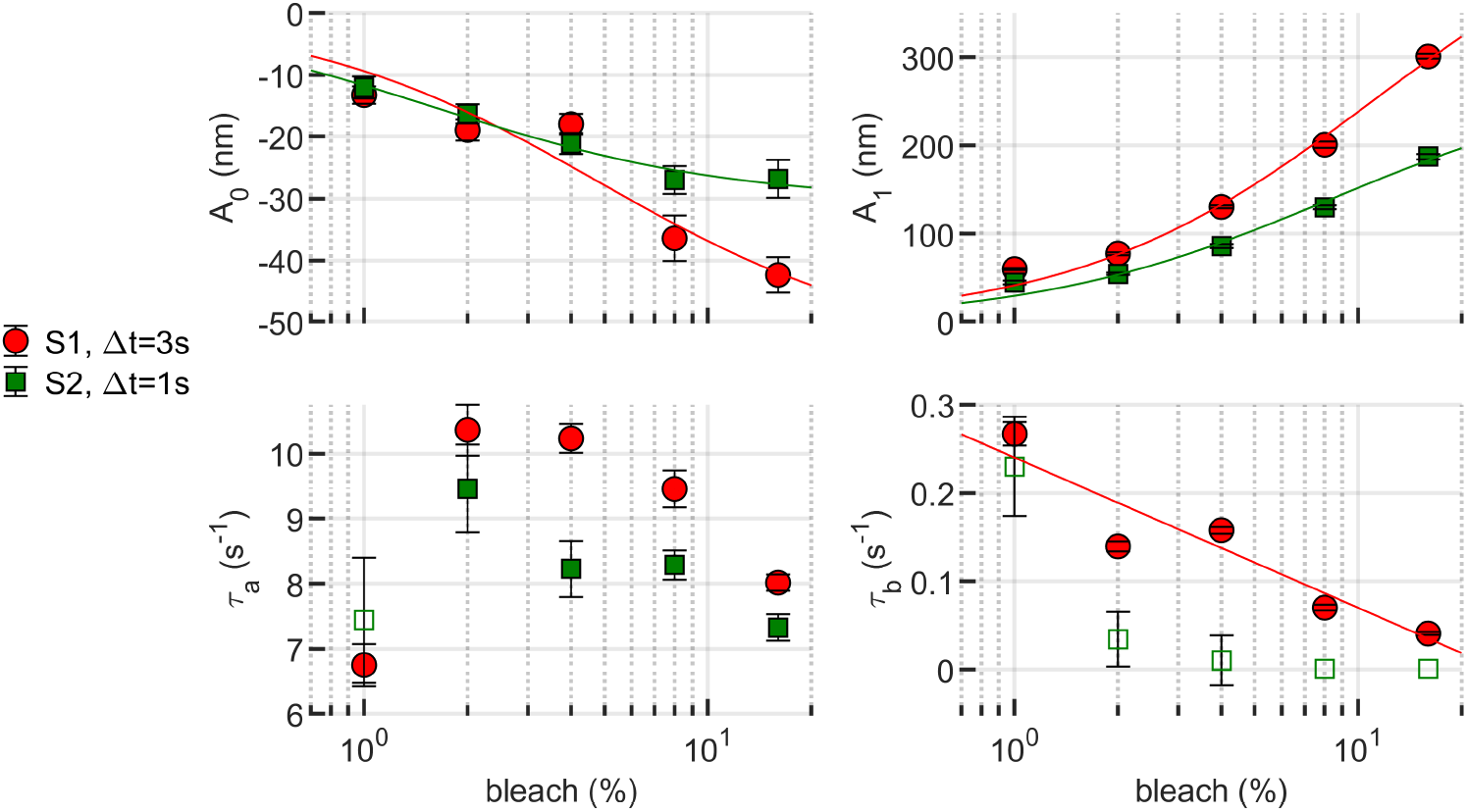
Fitting parameters as a function of bleaching level. *A*_0_, *A*_1_ and *τ*_*B*_ exhibit well-behaved patterns, either monotonically growing or decreasing. The qualitative observation that subject 1’s responses–both the contraction portion and the elongation portion–were stronger than subject 2’s is clearly visible in the plots of *A*_0_ and *A*_1_. Meanwhile, *τ*_*A*_ peaks at 2% bleaching for both subjects. *A*_0_ and *A*_1_ were fit with a sigmoidal Michaelis–Menten function, which fit both parameters in both subjects well (*R*^2^ ≥ 0.85). Based on the results shown in Supplementary Fig. 1, only the *τ*_*b*_ estimates from 3 s measurements from subject 1 were fit, this time with a log-linear model (*R*^2^ = 0.84). Estimates of *τ*_*b*_ from subject 2’s 1 s measurements are shown as unfilled markers. The error bars indicate the 95% confidence bounds of the fitting parameters.

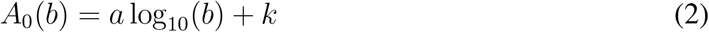

where *b* is the bleaching percentage and the free parameters are given by *a* = −25.1 nm and *k* = −10.7 nm for subject 1 and *a* = −13.4 nm and *k* = −12.6 nm for subject 2, with a goodness of fit of *R*^2^ = 0.84 and *R*^2^ = 0.93, respectively. This simple log-linear model is a good predictor of the early contraction stage within the experimental bleaching range but it has some undesirable properties outside of this range—at 0% and 100% bleaching, for instance. Taking that into account, the data was also fitted to a sigmoidal Michaelis–Menten equation (*15*):

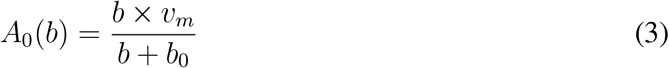

with free parameters of *v*_*m*_ = −54.6 nm and *b*_0_ = 4.8% and goodness of fit *R*^2^ = 0.85 for subject 1 and *v*_*m*_ = −30.5 nm and *b*_0_ = 1.6% and *R*^2^ = 0.97 for subject 2. These fits are at least as good as the log-linear fit above, and have the additional benefit of being well-behaved for bleaching percentage extrema.

The elongation amplitude term *A*_1_ exhibited a nonlinear, monotonically increasing relationship to dose. Fitting *A*_1_ to the Michaelis–Menten equation gives *v*_*m*_ = 502.3 nm and *b*_0_ = 11.1% with *R*^2^ = 0.98 for subject 1 and *v*_*m*_ = 279.5 nm and *b*_0_ = 8.4% with *R*^2^ = 0.97 for subject 2.

A curious result was obtained for elongation time constant *τ*_*a*_, with a peak around 2% bleaching for both subjects. This may be an artifact of the relatively low SNR of the 1% bleaching measurements, and subsequent underestimation of *τ*_*a*_. The question merits further investigation, and if the peak at 2% is reproducible, it’s a surprising and interesting finding.

Finally, *τ*_*b*_ was observed to decay monotonically for both subjects. Due to the diminished confidence in this estimate using a 1 s measurement duration (see Supplementary Fig. S1), the analysis was restricted to subject 1. Those fitted parameters displayed an apparent linear behavior on a semi-log scale, characterized by *a* = −0.17 nm and *k* = 0.24 nm, with *R*^2^ = 0.84. For completeness, the data for subject 2 was included in the plot, represented by unfilled markers.

Measurements were extended to higher bleaching levels (32% and 64%), as depicted in Supplementary Fig. S2. For these, the observation window was restricted to 1% for both subjects due to challenges in sustaining fixation after higher bleaching doses. Saturation in amplitude was not observed in either subject. Interestingly, the fitting diverges from the proposed model at these levels, which could indicate the presence of an additional exponentially-rising, lower-amplitude component in response to higher energy stimuli, as discussed in (*17*). Alternative hypotheses include a divergence from linear behavior due to nonlinear biomechanical factors (e.g., drag or other hydrodynamic factors). However, the lack of data on the recovery portion of ORGs at these bleaching levels limits our ability to model these factors.

### 2.2 Paired flashes

In both subjects we measured responses to two successive flashes, each lasting 10 ms and having a power of 5.05 µW (equivalent to 4% single-flash bleaching). The flashes were separated by an inter-stimulus interval (ISI) denoted as *t*_isi_, ranging from 15 ms to 300 ms.

Our observations revealed a cumulative, nonlinear effect when two equal flashes were presented. The cumulative response to two flashes that each bleach 4% of photopigment is equal in maximum excursion (~150 nm) to a single flash bleaching 8% of photopigment, but smaller than twice the ~100 nm response to a 4% bleach. The response to two 4% flashes is shown in relation to single flash responses in Fig. 4 (left). Flashes whose onsets were separated by as little as 15 ms result in distinguishable OS contractions, as seen in Fig. 4 (right).

**Figure 4.**
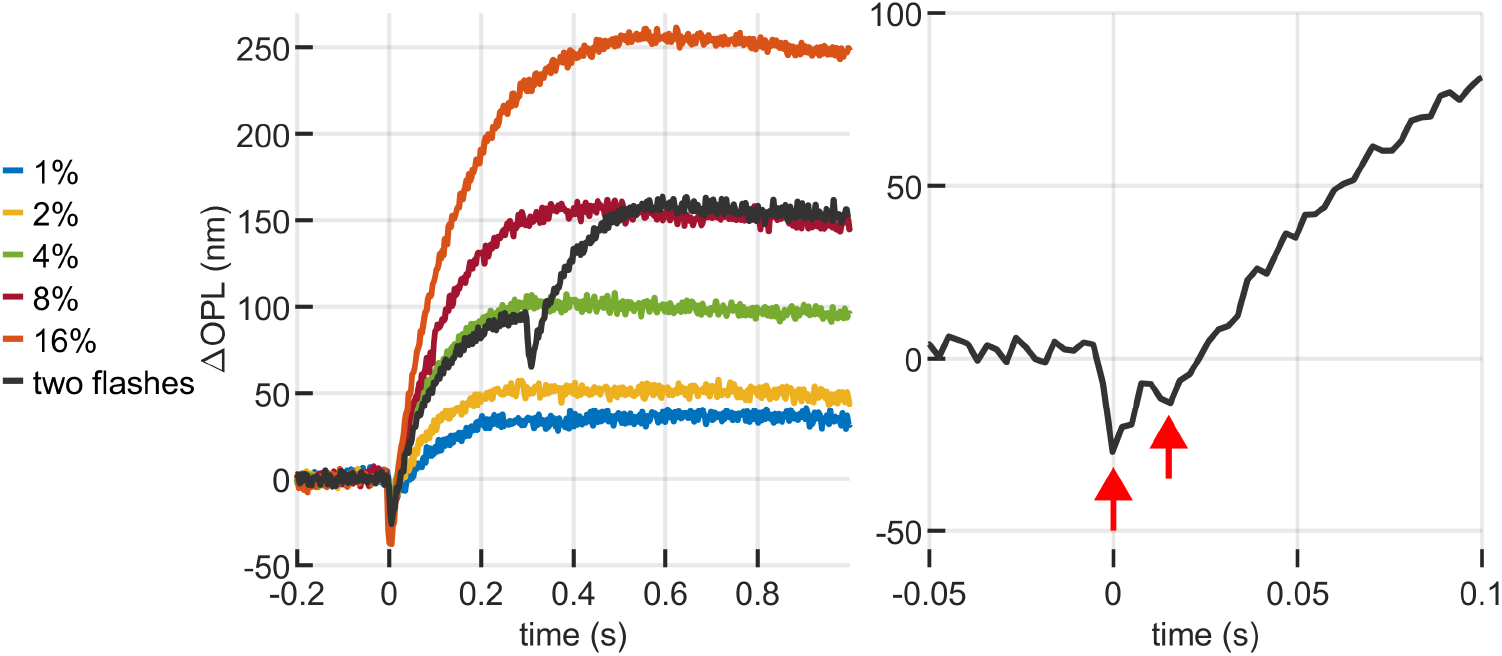
Responses to paired flashes. (left) The response to two sequential flashes of 4% bleaching exhibits a nonlinear cumulative effect, with maximum elongation equal to a single flash of 8% bleaching; (right) Flashes as close as 15 ms apart can still be distinguished at the early response contraction.

The responses of cones to paired flashes had these characteristics over a range of interstimulus intervals between 15 ms and 300 ms, as seen in Fig. 5. Raw data are plotted with colored markers; fits to the shifted-sum, two-flash model described in Eq. 14 are plotted with a solid black line. Distinct contractile responses are visible for all first and second flashes (although the 15 ms ISI response is more visible in the zoomed view in Fig. 4 (right)). Qualitatively, it can be seen that the magnitude of the second OS contraction appears to increase with increasing inter-stimulus interval. It also appears as though the elongation accompanying the second flash is attenuated when compared with the first. For example, in the *t*_isi_ = 300 ms response shown in Fig. 5 (right), the first flash appears to elongate the OS by ~100 nm, and the second by ~50 nm.

**Figure 5.**
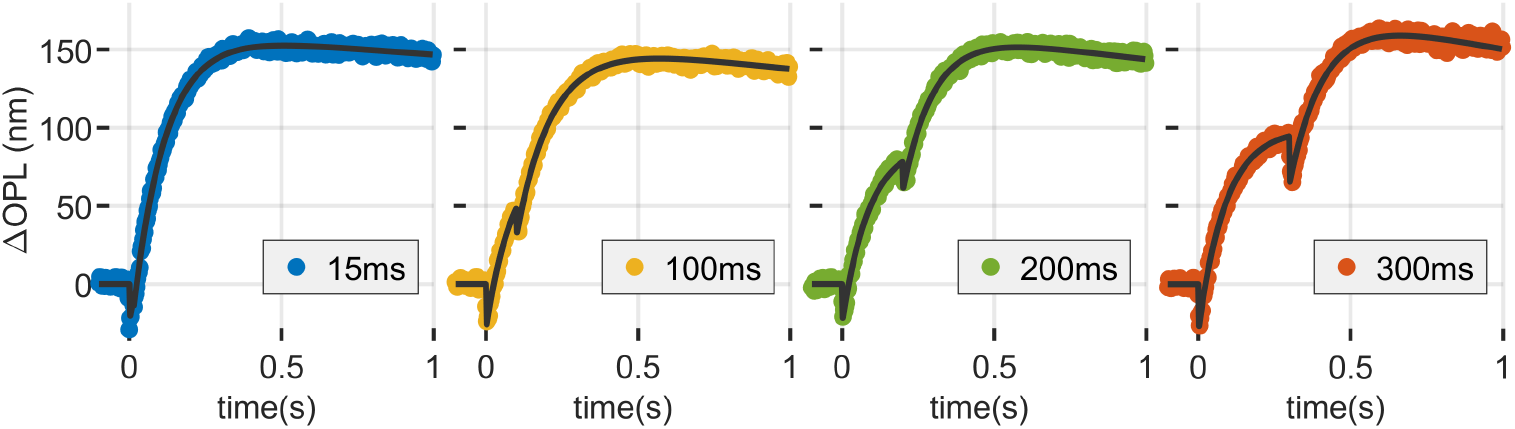
Responses to pairs of flashes separated by *t*_isi_ of 15, 100, 200 and 300 ms. Responses to two 4% flashes results in a total OS elongation of ~150 nm, equivalent to the elongation generated by a single 8% flash. However, the responses to multiple flashes are not simply the sums of individual responses. In the case of contraction, this can be seen in the magnitude of the early contraction, which seems to increase with increasing *t*_isi_. In the case of elongation, the nonlinear additivity is most apparent in the *t*_isi_ = 300 ms case (right), where the elongation caused by the second flash appears attenuated compared to that caused by the first. The maximum elongation for all four conditions was the same, and the overall amplitude of response of the second flash was reduced by the first.

Fitting with the shifted-sum, two-flash model yielded estimates of eight free parameters, four for each of the two responses, labeled *α* and *β* for clarity: contraction amplitudes *A*_0,*α*_ and *A*_0,*β*_; elongation amplitudes *A*_1,*α*_ and *A*_1,*β*_; elongation time constants *τ*_*a*,*α*_ and *τ*_*a*,*β*_; and recovery time constants *τ*_*b*,*α*_ and *τ*_*b*,*β*_. For each of these pairs, we sought to understand the relationship between the first and second response’s parameters, i.e., the effect of the first parameter on the second. To assess this, we visualized the ratios of the *β* to *α* estimates as functions of inter-stimulus interval *t*_isi_. These ratios are shown in Fig. 6.

**Figure 6.**
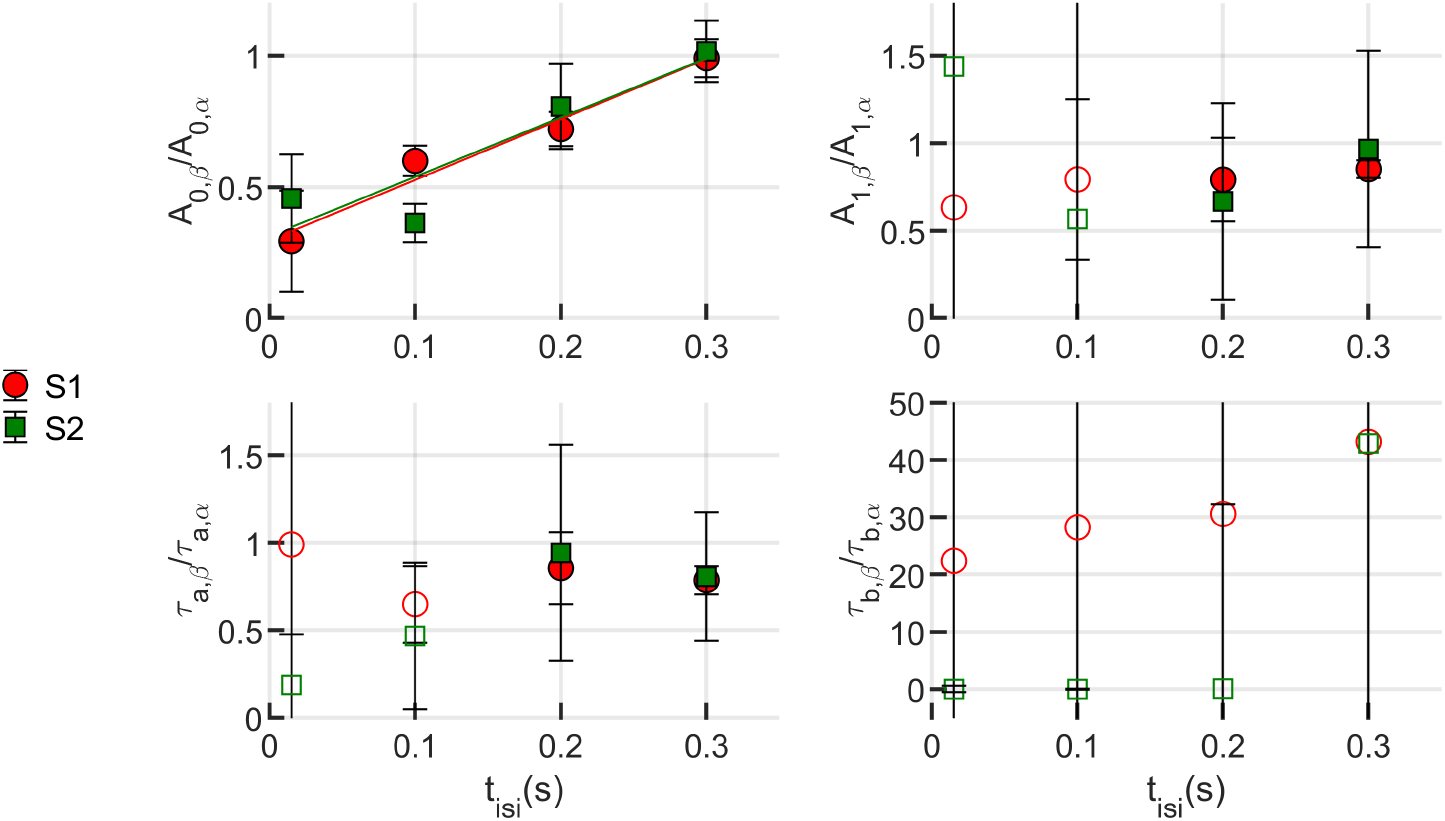
Responses to paired flashes. The impact of the first flash on the second flash response was quantified by fitting the total response with the shifted-sum model shown in Eq. 14. From this fit, four free parameters could be estimated to describe the parts of the response due to each of the flashes, and the ratio of the second estimate to the first could be visualized as a function of *t*_isi_. For *A*_0_, a clear relationship with *t*_isi_ can be seen, with longer *t*_isi_ associated with higher *A*_0_ magnitude. Differences between the *A*_1_ ratio for *t*_isi_ = 200 ms and *t*_isi_ = 200 ms are visible, but smaller than the error bars shown. For some parameters the observation window of the first flash was too small, leading to unreliable estimates. Those values are plotted with unfilled markers. All the values of *τ*_*B*_ falls into this category and are only shown here for completeness. Error bars were calculated by propagation of uncertainty of the confidence bounds of the fitting parameters.

Trust in the fitting-based estimates of response parameters depended on the duration of measurement, as shown in Supplementary Fig. S1. Here, the inter-stimulus interval and total recording time both imposed limits on the number of samples used to estimate a parameter.

Because the initial contraction is rapid, it can be estimated well with relatively few samples. The elongation amplitude *A*_1_ and time constant *τ*_*a*_ require relatively more than 100 ms to estimate reliably, thus we rely solely on the 200 ms and 300 ms ISI estimates. For intervals *t*_isi_ ≤ 100 ms, we lack confidence in the accuracy of the *A*_1_ and *τ*_*a*_ estimates. Because these measurements were all collected within one second, we are also tentative about estimates of *τ*_*b*_ for all *t*_isi_. The tentative estimates of *A*_1_, *τ*_*a*_, and *τ*_*b*_ are plotted, for completeness, with unfilled markers.

The ratio *A*_0,*β*_*/A*_0,*α*_ (Fig. 6, upper left) bears a clear relationship to *t*_isi_, with *A*_0,*β*_ being attenuated in inverse proportion to *t*_isi_. For the shortest inter-stimulus interval *t*_isi_ = 15 ms, the second contraction *A*_0,*β*_ is less than half the magnitude of *A*_0,*α*_, despite the first flash having bleached only 4% of photopigment. The slope of a linear fit to the ratios was 2.30 and 2.26 for subjects 1 and 2, respectively, and for both subjects a ratio of 1 was reached when *t*_isi_ ≈ 0.3 s.

For *t*_isi_ ≥ 200 ms, fitting of *A*_1_ and *τ*_*a*_ resulted in acceptably low fitting error. For both of these parameters, the ratio of second to first response was between 0.75 and 1, as shown in Fig. 6 (upper right and lower left, respectively). Unfortunately, because of the low reliability of the 15 ms and 100 ms fits, a trend is difficult to establish, but we will later consider the cases where these ratios are constant with respect to *t*_isi_ and the cases where they are correlated or anticorrelated with *t*_isi_, respectively.

To address uncertainty, error bars on the graph are computed by propagating the uncertainty arising from the confidence bounds obtained during the fitting process. Throughout the fitting procedure, the level of certainty of the free parameters were set as 95%, establishing lower and upper bounds. The width of this interval indicates the degree of uncertainty regarding the fitted coefficients. With that, the uncertainty of the ratio of given fitting parameters, for instance *A*_0,*α*_ and *A*_0,*β*_, was determined using the variance formula for error propagation, assuming independence among the variables.

### 2.3 Stimulus pulse train

The attenuation of the ORG response due to the influence of a preceding flash, observed above, was corroborated through a sequence of multiple flashes. For this purpose, we recorded cone responses to a series of 1 ms flashes, each bleaching 4% of photopigment and separated by an interval of 50 ms. A distinct contraction was observed after the first few flashes, gradually diminishing in magnitude until it became indistinguishable from the background signal noise. This pattern of diminishing response aligns with the observations made during the paired flash stimulus experiments. The results of this experiment are shown in Fig. 7. No effort was made to fit these using the model because the *t*_isi_ of 50 ms does not permit reliable fitting of the responses.

**Figure 7.**
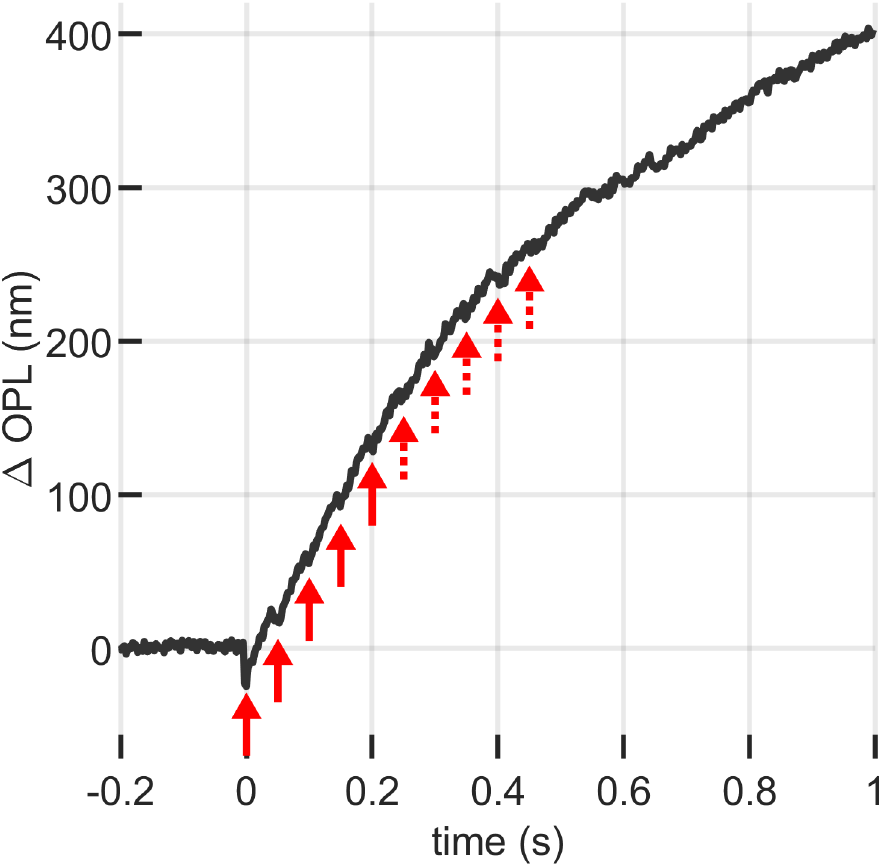
ORG response to a series of 10 flashes of 10 ms, 4% bleaching flashes separated by 50 ms. It can be seen that the early stage contraction is increasingly attenuated by the presence of the previous flash (solid line arrows) until it gets hidden by the noise (dashed arrows).

### 2.4 Adapting background

In these trials, following a 5-minute period of dark adaptation, the studied region of the retina is exposed to a low-intensity background light for a duration of 10 seconds before data acquisition begins. Subsequently, a single 10 ms flash with a photobleaching efficiency of 4% is administered.

The presence of the low-intensity background light had a noticeable impact on the overall response dynamics (Fig. 8). It was observed that this background light caused sustained, apparently constant-rate OS elongation, to which the flash response apparently added. The response was thus modeled as the sum of a linear component and the single flash response (Eq. 1):

**Figure 8.**
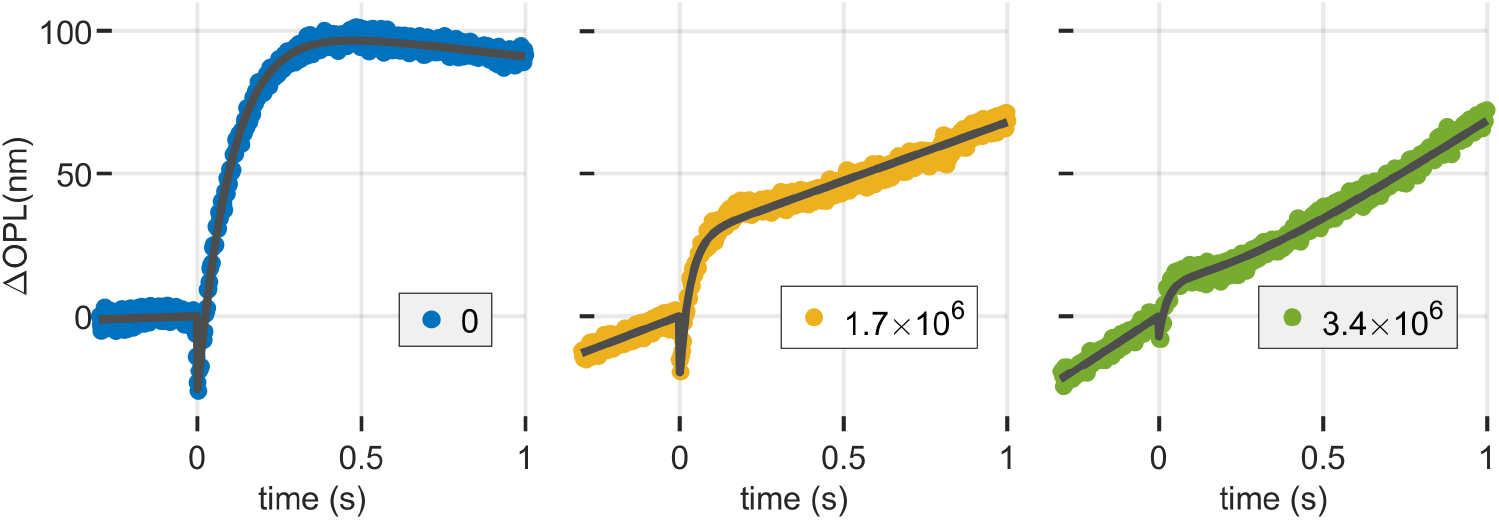
Optoretinograms of a 4% flash in the presence of a dim background light using the same 555 nm source. Three powers, measured at the cornea, were used: 0 nW, 114 nW, and 236 nW, which correspond to photon flux densities of 0 s^−1^µm^−2^, 3.1 × 10^6^ s^−1^µm^−2^, and 6.5 × 10^6^ s^−1^µm^−2^. After adapting for 10 s to the background light, the ORG presented a slope superposed to the response of the flash released at *t* = 0. The presence of background also caused a reduction in both the contraction and elongation components of the cones’ response to the flash, in spite of bleaching relatively small amounts of photopigment.

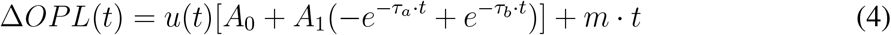

The resultant fitted parameters, as a function of background illumination, are shown in Fig. 9. It was observed that the slope (*m*) of the elongation due to the dim background line was directly proportional to the background light intensity, with proportionality constants given by 11.97 nm^3^*/*photons and 8.85 nm^3^*/*photons for subjects 1 and 2 respectively.

**Figure 9.**
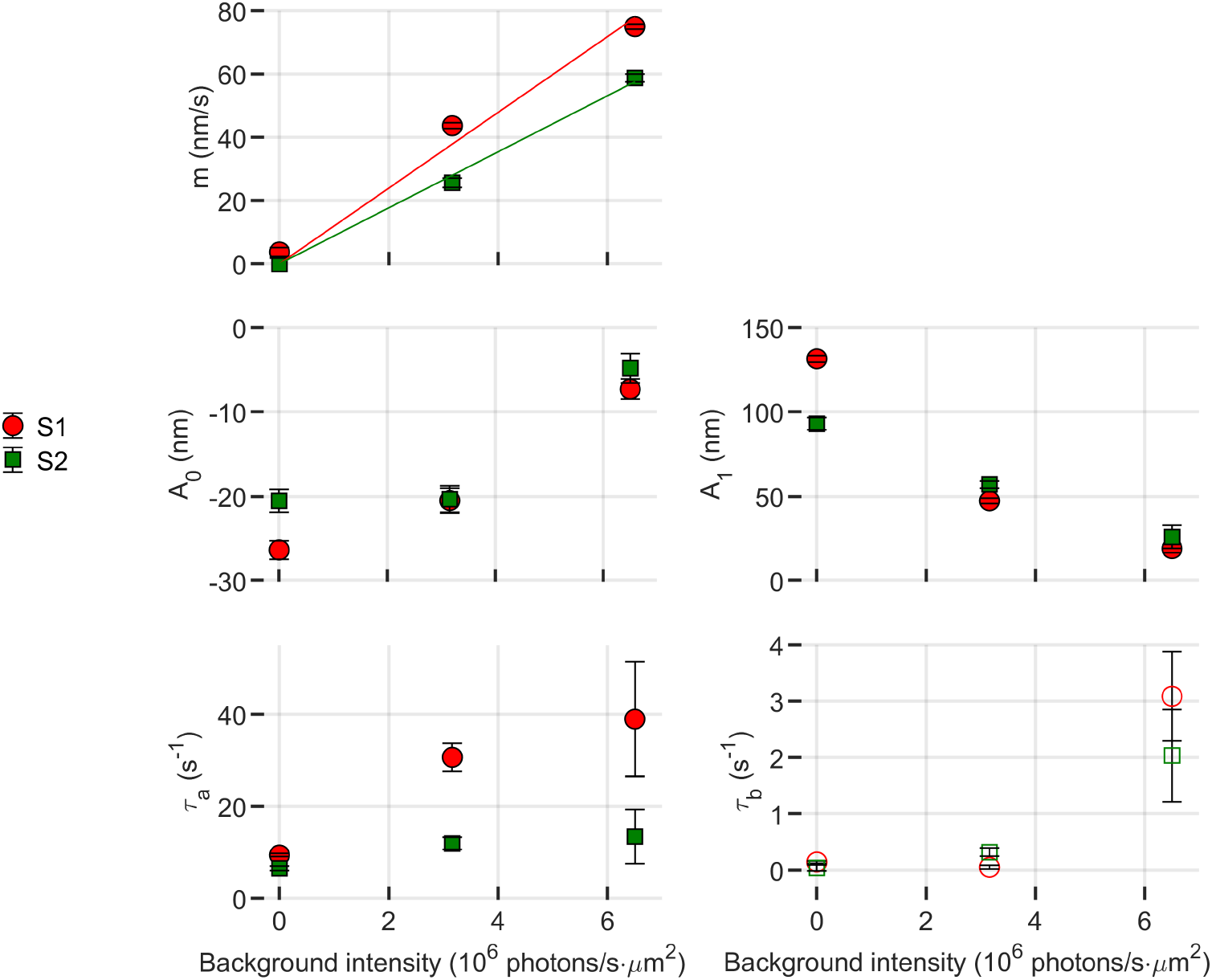
Fitting parameters of Eq. 15 as a function of background illumination level. The constant elongation rate (*m*) appears to increase linearly with illuminance. The magnitudes of the contraction (*A*_0_) and elongation (*A*_1_) components of the response appear to be attenuated in the presence of a background.

These backgrounds bleached 0%, 60%, and 85% of photopigment in the 10 s prior to the stimulus flash. It is apparent that at the two higher bleaching levels parameters *A*_0_, *A*_1_, and *τ*_*a*_ are affected by the prior bleaching, as shown in Fig. 8. The magnitudes of *A*_0_ and *A*_1_ display a monotonic decay, with an apparent parabolic trend in relation to background intensity. The elongation rate *τ*_*A*_ shows an increasing trend with background intensity, closely resembling linear behavior.

*τ*_*B*_ falls within the minimum confidence window and is represented, for the sake of completeness, as unfilled markers.

## 3 Discussion

We have reported stimulus-evoked cone photoreceptor responses measured with adaptive optics (AO) full-field (FF) swept-source (SS) OCT. AO-FF-SS-OCT joins a growing body of imaging modalities to have reported ORG responses, including common-path interferometric methods (*9, 11*), FF-SS-OCT with digital aberration correction (*10*), point-scanning AO-OCT (*12*,*13*), line-scanning AO-OCT (*18*), and conventional scanning OCT (*16*). This methodological diversity is a boon to the nascent field of optoretinography, as each presents unique advantages and will likely be the optimal choice for some problem domains. However, it presents a challenge as well, since data produced by different methods are not inherently commensurate. Ultimately we need to have quantitative methods for harmonizing measurements across these methods. Such harmonization is especially critical if optoretinography has commercial potential, as it may preempt the analogous difficulties faced by researchers aggregating data from multiple OCT and OCTA instruments.

A critical step toward creating ORG standards is the development of methods to quantify photoreceptor responses. We have proposed such methods in previous publications (*12, 15, 16*), but the latter have been *ad hoc*, proposed in response to informal, visual inspection of the responses. These figures of merit (e.g., maximum OS excursion or OS elongation rate) may be intuitive, but lack sufficient formal specificity to be of use to other ORG investigators. The problem is worse for detection schemes other than position-based, phase-sensitive OCT. For example, it is not obvious how to translate the latter’s relatively transparent signal into the produced by phase-velocity based methods (*16*), *en face* methods (*9, 11*), or statistical approaches (*19*). On the other hand, a general model of photoreceptor responses may be used, in conjuction with knowledge of the imaging system and signal processing techniques.

A good ORG model will be sufficiently complex (measured in terms, e.g., of order and dimensionality) as to represent distinct and possibly uncorrelated aspects of the response, but sufficiently simple to avoid overfitting and generation of meaningless dimensions. Based on our visual inspection of the ORG responses reported here and previously by our group, we felt tha the model expressed by Eq. 1 strikes this balance. The model, based on an overdamped harmonic oscillator with an offset, does not aim to delineate the intricate biomechanical intricacies of the process but, rather, describes the mechanical axial change observed in the cone outer segment. With only four parameters, the fitting equation allows direct access to key intuitive aspects of the ORG response: amplitudes of contraction and elongation, and rates of elongation and recovery to baseline.

The purpose of this paper is not to propose a perfect solution to the problem of modeling photoreceptor ORG responses, but rather to get the ball rolling on such a project. Additional measurements by us or other groups could lead to refinements, revisions, or outright rejection of this model.

### 3.1 Single flash responses

As shown in Supplementary Table S1, the residual fitting error across all bleaching levels was high (*R*^2^ > 0.95 for all measurements, and *R*^2^ > 0.99 for most), which confirms the visual goodness of fit shown in Figs. 1 and 2. The goodness of fit was high even in the case of high (≥32%) bleaching levels, where the qualitative appearance of the fit (Supplementary Fig. S2) was not as good.

The model described by Eq. 1 appears qualitatively to capture the main features of the ORG response, as illustrated by Figs. 1 and 2. Fig. 1 panels (c) and (d) illustrate the role of each of the model’s free parameters on the pattern of OS deformation. Hypothetically, these components of the response may be tied to electrostatic effects of photoisomerization (*20*) (*A*_0_), total osmolar transit and biomechanical constraints (*A*_1_), osmolar gradients (*17, 21*) (*τ*_*a*_), and phototransductive downregulation or photopigment regeneration (*τ*_*b*_). Dependence of these aspects of the response on experimental parameters such as stimulus characteristics, eccentricity, and outer segment length would form the basis of a normative database. Disease-related deviations from the resulting norms could potentially then be expressed in terms of the model parameters, which in conjuction with the clinical work-up could lead to specific, testable hypotheses about disease mechanisms. In return, when disease etiology is well-understood, such deviations could supply natural experiments to further elucidate the mechanisms of the ORG.

### 3.2 Potential standardization with alternative ORG metrics and implementations

As described above, interoperability of ORG methods is a key motivation for this work. As such, in order to compare with previous works, we derived a few figures of merit from the main model. *Ad hoc* figures of merit that have been reported include the early elongation slope *m*_0_, time to maximum excursion *t*_*max*_, and maximum elongation Δ*OPL*_*max*_. These intuitive figures of merit can be derived from the derivative of Δ*OPL*(*t*):

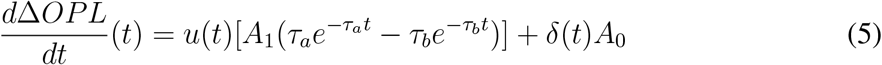

While rate of early elongation is not well-defined, the elongation rate can be calculated at any desired time using Eq. 5. The root of Eq. 5 gives expressions for *t*_*max*_ (Eq. 6) and Δ*OPL*_*max*_ (Eq. 7), the time at which maximum outer segment elongation is attained, and the Δ*OPL* at that time, respectively:

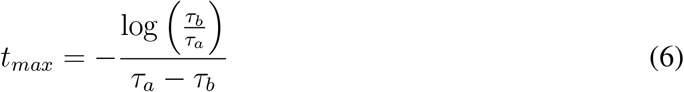

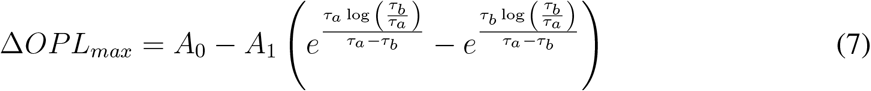

The data collection was confined to only two volunteers, a decision driven by the substantial size of each generated file and the extensive time required for data processing. While this approach is useful for measuring the response of individual cones, it restricts the feasibility of large-scale data acquisition, crucial not only for establishing normative behavior of healthy retinae but also for offering clinical insight into pathologies. Another limitation is the high cost associated with OCT with enough speed and resolution, capable of acquiring ORGs, which restricts the number of groups able to develop such a system.

Alternative implementations include a method based on OCT phase velocity, where the velocity of retinal structures is monitored rather than their position (*16*). Planned future work includes reproduction of some of these results in a larger number of subjects using the velocity-based approach. This method allows measurement of cone responses without the need to track specific cells over time, thus dispensing with the needs for an adaptive optics subsystem, digital aberration correction, real-time tracking, and three-dimensional segmentation and registration. Moreover, because the velocity and position of the OS reflectors are related by integration, there is potential to use the derivative of Eq. 1, shown in Eq. 5.

The evaluation of human cone photoreceptor function can also be achieved through 2D imaging techniques such as a fundus camera (*9*) and scanning light ophthalmoscope (SLO) (*11*). In these methods, there is no direct assessment of outer segment length, and functional responses are encoded in the individual cone photoreceptor reflectance. This change in brightness is hypothesized to be related to constructive and destructive interference from the light reflected at the photoreceptor inner segment - outer segment junction (ISOS) and cone outer segment tip (COST), thereby establishing a correspondence with OCT measurements (*22*). While the 2D approach doesn’t directly capture morphological changes, it offers the advantages of more affordable setups and significantly smaller volumes of data per ORG acquisition. The model proposed in this work could also be used to model cone responses measured in *en face*, commonpath interferometry, using an equation such as:

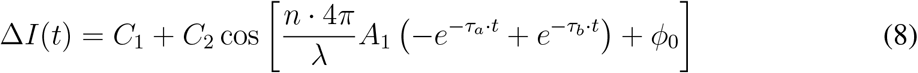

The proposed model for light-evoked outer segment (OS) deformations could potentially serve as a valuable tool for directly comparing observed results across various OCT setups for ORG and potentially to act as a bridge linking 2D measurements with the optical path length variation measured using OCT.

### 3.3 Alternative models

In addition to the model described by Eq. 1, a number of other models were considered. These are listed in Supplementary Table S2 and visually dipicted in Fig. 10. For clarity, we refer to them by the colors of the plots in Fig. 10. The model used in the results above was the magenta model (Eq. 1). It is worth noting that as *t* → ∞, the model converges to *A*_0_, resulting in negative values. To address this, the green model 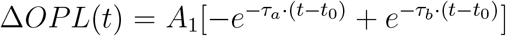 was proposed, which converges to Δ*OPL* = 0 as *t* → ∞. Over the first seconds of observation, the behaviors of the magenta and green models are very similar. In Fig. 10, the two visually overlap, and were offset by ±1 nm to facilitate visualization.

**Figure 10.**
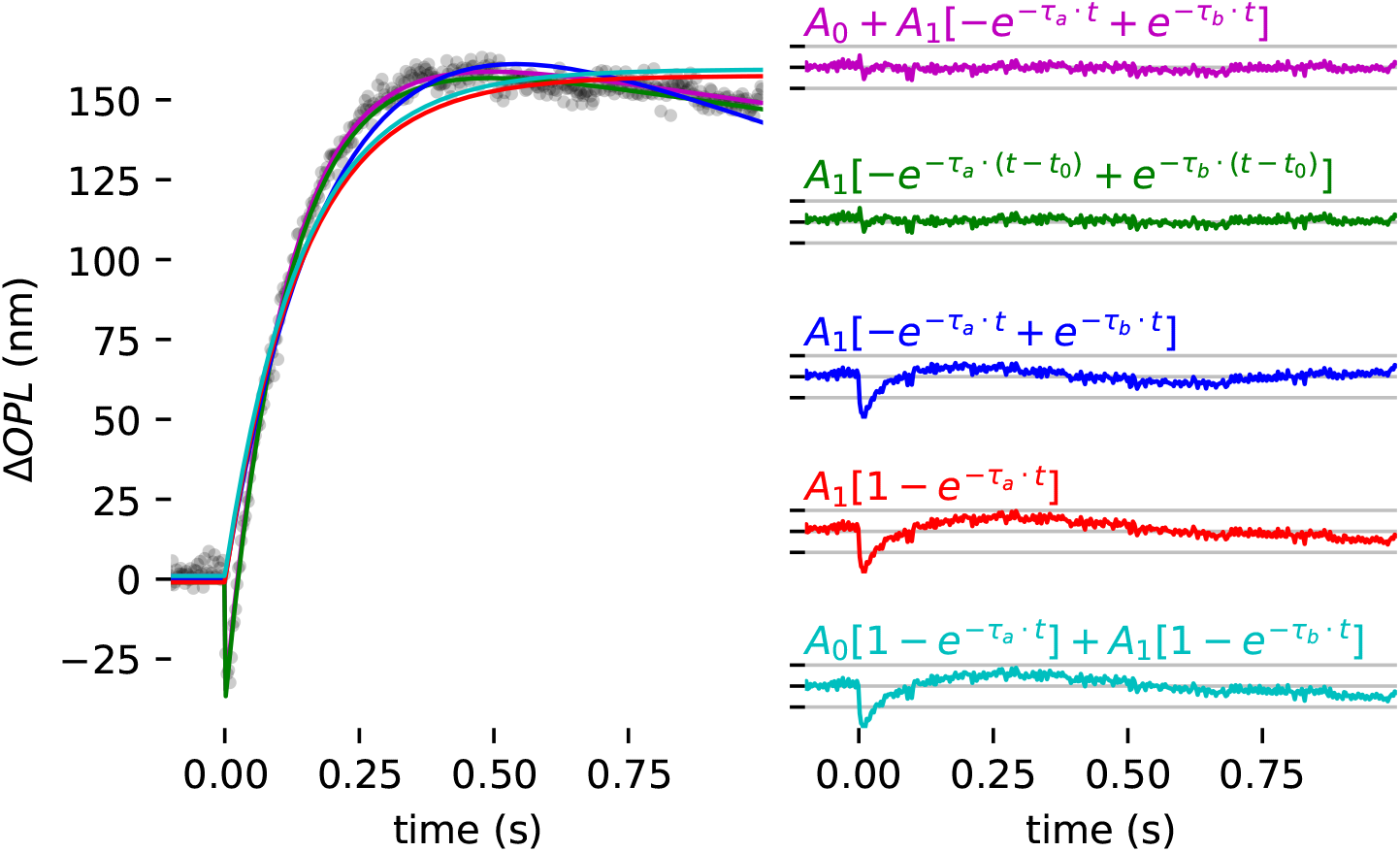
Comparison among models of residual fitting error. (left) Experimental data fitted for different models (left) and corresponding residual fitting error for each model (right).

The red and blue models represent our earliest attempts to quantify the cone responses, and are described here for the sake of completeness. While the red might be of interest of capturing the osmotic elongation, it does not predict the early contraction or recovery to baseline. Mean-while, the blue model takes into account both elongation and recovery and may be of interest to investigators using slower modalities (e.g., conventional OCT or AO-SLO). Those two models are simpler than the others and may better prevent overfitting of data. On the other hand, they potentially provide less rich sets of biomarkers of the cone response. The cyan model was in-spired by the recent findings of Pandiyan et al. (*17*), which showed that the cone responses they measured–particularly those to the highest energy stimulus flashes–were better fit by a pair of exponential functions with different amplitudes and time constants.

To compare the models, they were all used to fit an exemplary cone response to an 8% bleach. The magenta and green models yielded the lowest residual errors, which were nearly equivalent (differing by less than 0.001 nm). To facilitate visualization, they are offset vertically in Fig. 10 by ±1 nm. This was also true of the cyan and red models, which had significantly higher residual error than the magenta and green, and they were similarly offset in Fig. 10.

We did not find the cyan model 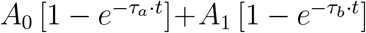 to perform better than the red model 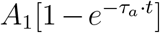. In unconstrained, unbounded fits, the cyan model produced roughly equal time constants *τ*_*a*_ and *τ*_*b*_ (similar to the estimate of *τ*_*a*_ produced by fitting the red model) and roughly equal amplitudes *A*_0_ and *A*_1_ (each approximately half of the *A*_1_ obtained by fitting the red model). Even at the brightest bleaching levels we used (32% and 64%, where the magenta and green models don’t appear to fit perfectly (see Supplementary Fig. S2), the cyan model’s residual error was higher than that of the magenta or green models. The red and cyan models were so similar in appearance that they were also offset vertically by ±1 nm to make they both visible.

The blue, red, and cyan models are not designed to capture the initial contraction of the OS. To fairly facilitate comparison among those, Supplementary Table S2 lists separate residual RMS error for the entire duration (−0.1 ≤ *t* ≤ 1.0 s) and the residual RMS error for *t* ≥ 0.5 s.

While the magenta and green models are able to capture the magnitude of the initial negative Δ*OPL*, they are still too simple to model the rate of contraction of the early response. This simplification was by design, as our volume rate of 400 Hz was too slow to collect multiple measurements during the cone OS contraction. To comprehensively model this specific portion of the overall response of the ORG, a considerably faster acquisition rate would be imperative.

While not identical, the magenta and green models exhibit similar performance within an observational window of a few seconds, both offering insights into the early OS contraction. The magenta model fits the amplitude of the early stage contraction, while the green model fits its duration. Both pieces of information can be derived from either model. We have opted to highlight amplitude, a parameter that we felt to be a more obvious or intuitive descriptor of the response. Future work may show whether one equation proves more advantageous in identifying overall ORG response mechanisms or characterizing disease-related ORG changes.

Both models exhibit limitations in characterizing the dynamics of a single flash with high doses, failing to accurately reflect the experimental data in both the early contraction and elongation phases. These deviations suggest that additional components may be necessary to fit responses to high-energy stimuli. It is possible that could be additional exponential components, as proposed by Pandiyan *et al*. (*17*), or perhaps other nonlinear components related to hydrostatic or biomechanical constraints (e.g. drag between the OS and its surroundings, or trans-membrane pressure inequalities).

### 3.4 Complex stimuli

The outer segment of human cones exhibits a distinct mechanical response when subjected to bleaching stimuli. Following a brief, single flash, this reaction manifests as a rapid contraction succeeded by a gradual elongation and an even slower return to baseline. The magnitude and rate of these processes are tied to the dosage of the stimulus, and the overall dynamics can be described as the sum of two exponentials with identical amplitudes, along with a constant (Eq. 1).

Other functions, derived from this model, were investigated to analyze more intricate stimulus patterns, such as multiple flashes and a flash in the presence of a dim background. Through this exploration, it became evident that the response of the photoreceptor–including both the initial contraction and the subsequent elongation–is attenuated in the presence of a preceding stimulus, even when the stimulus is significantly below the expected cone pigment saturation level.

Results of the two-flash experiment revealed that the maximum excursion following the second flash remained unaltered, regardless of the interval between flashes. This observation implies a sole dependency on the cumulative bleaching level, hinting at a potential correlation between elongation and photopigment availability.

Meanwhile, the attenuation of the amplitude of contraction appeared to be inversely proportional to the inter-stimulus interval. It reached a relative amplitude of 1 at around 300 ms, close to *t*_*max*_. Notably, the attenuation of the second contraction exhibited remarkable consistency across both subjects. The attenuation of the overall ORG response in the presence of a preceding stimulus not only allows further discussion about the mechanisms at play in each phase of the ORG response but also highlights the critical importance of a consistent dark adaptation routine.

### 3.5 Conclusion

By allowing noninvasive, objective measurement of neural function in the retina, with the ability to localize functional responses and precisely observe structure-function correlations, the ORG offers novel, complementary OCT-based biomarkers of functional responses of neurons and other structures. As such, it has the potential to transform clinical assessment of retinal disease and accelerate the development of novel therapeutics.

The presented model for light-evoked outer segment (OS) deformations offers a comprehensive understanding of the mechanical response observed in human cone photoreceptors following bleaching stimuli. This model, with only four parameters, allows direct access to crucial aspects of optoretinography (ORG) responses, including the amplitudes of early contraction and later elongation, as well as the rates of elongation and recovery to baseline. The model fits well to the experimental data with a variety of stimulus intensity and patterns. Notably, the experimental data supporting this model were collected from only two subjects, underscoring the need for future studies with larger cohorts to establish a more generalized understanding of the average behavior in healthy retinas and its implications in pathological conditions.

**Supplementary Material** accompanies this paper at http://www.scienceadvances.org/.

## 4 Materials and Methods

### 4.1 Imaging system

The imaging system has been reported in detail elsewhere (*23*). A schematic of the system is shown in Supplementary Fig. S3. Briefly, it consists of a Mach-Zehnder interferometer with a tunable light source (825 nm to 875 nm, BS-840-2-HP, Superlum, Cork, Ireland) divided into sample and reference arms by a polarizing beamsplitter (PBS). The reflected light goes to the sample arm and the transmitted is expanded and collimated, striking the camera at an angle of ~1° with respect to the sample arm in order to create a spatially-varying phase delay between the sample and reference fields and thus a carrier frequency. The resulting modulation of acquired interferograms allows the filtering in Fourier space of the retinal interference signal from DC components and common-path coherent artifacts (*24*), ensuring that the demodulated interferograms will consist only of signal generated by interference between the reference mirror and sample.

In the sample arm, light illuminated a 2° field-of-view (FOV) on the retina with a converging (but not focused) beam with a power of 3 mW measured at the cornea. Ocular aberrations were corrected in real-time by an adaptive optics (AO) subsystem operating in a closed-loop using open-source software developed in Python/Cython by our lab (*25*). By measuring and correcting aberrations over a 6.75 mm diameter pupil, it provided a diffraction-limited lateral resolution of 2.6 µm in the retina. A superluminescent diode (SLD, 755 nm, 30 nm FWHM) was used as AO beacon (IPSDM0701-C, Inphenix, Livermore CA, USA).

Sample and reference arms are recombined onto a high-speed 2D CMOS sensor (FAST-CAM NOVA S-12, Photron, Tokyo, Japan) running at 200 kHz. The spectral sweep was sampled with 500 frames from the camera, which resulted in an OCT volume rate of 400 Hz. The camera’s pixels are 20 µm wide, and magnification between the retina and camera was 22× (assuming a 16.7 mm focal length for the eye), and thus each pixel sampled ~0.9 µm in the lateral dimensions. The source swept over 50 nm of bandwidth between 875 nm and 825 nm, resulting in axial resolutions of 6.4 µm and 4.6 µm in the air and eye (*n* = 1.38), respectively. The retinal depth was sampled with an interval of 2.3 µm or frequency of 435 mm^−1^.

### 4.2 Stimulus delivery and characterization

The setup also incorporated a visible-light channel with precise control of flash time and power, designed for stimulating the retina. A fiber-coupled 565 nm light emitting diode (M565F3, with DC4100 four-channel LED driver, Thorlabs; Newton, NJ, USA) combined with a 23 nm bandpass filter centered at 555 nm (FF01-554/23; Semrock; Lake Forest, IL, USA) was used as light source. At this wavelength, L and M cones are bleached equally. The illuminated area was limited to a circle with ∅ = 360 µm due to the low power produced by the bandpass filtered LED.

The stimulus pattern, delay, and duration were controlled using a function generator (Rigol DG4202, Suzhou, China), triggered by the CMOS camera. For a single flash, the resulting bleaching levels and corresponding optical powers and pulse duration are listed in Supplementary Table S3. Values correspond to an assumed ocular transmission of 1.0 and circular stimulated area with diameter 360 µm. Pulse width was 10 ms for all but the highest bleaching level of 64% (in the bottom row), for which the pulse width was broadened to 12.6 ms.

### 4.3 Human subject imaging

Two subjects, free of known retinal disease, dark-adapted, dilated and cyclopleged using topical drops of phenylephrine (2.5%) and tropicamide (1.0%), were imaged at 2° temporal to the fovea. A bite bar and a forehead rest were employed to position and stabilize the subject’s pupil during imaging while a calibrated target guided the subject’s fixation.

Subjects underwent a five-minute dark adaptation period before the imaging procedure. This involved placing the subject in a darkened room and covering the eye to be imaged with a patch. After the five-minute dark adaptation period, and just before commencing the imaging process, the subject was briefly exposed to ambient light within the darkened room for a few seconds to permit fixation and closing of the AO loop. OCT volumes were then acquired at a rate of 400 volumes per second for periods of 1 s or 3 s, with the stimulus flash delivered after 200 ms.

All procedures were in accordance with the Declaration of Helsinki and approved by the University of California Davis Institutional Review Board. The simultaneous illumination from the three sources was in accordance with laser safety standards (*26*).

### 4.4 Stimulus patterns

Development of the ORG model described below was based on measurements using a single flash of the stimulus source. However, to demonstrate the feasibility of using more complex stimuli, additional patterns were used:

1. *Single flash*. In these trials, stimuli were delivered at varying bleaching levels. The flash duration was set at 10 ms for most trials, except for the 64% trial, for which the duration had to be extended to 12.6 ms to compensate the low power output provided by the bleaching light source.
2. *Paired flashes*. In these trials, the subject received a pair of 4% flashes, separated by in inter-stimulus interval (*t*_isi_) of 15 ms, 100 ms, 200 ms, or 300 ms.
3. *Pulse trains*. In these trials, a sequence of 1% flashes were administered with an interval of 50 ms between each flash. The first flash was initiated 200 ms after the commencement of data acquisition.
4. *Adapting background*. In these trials, single 4% flashes were delivered against an adapting background. The subject were given a 10 s period to adapt to the background before the flash was delivered. For adaptation, three powers, measured at the cornea, were used: 0 nW, 114 nW, and 236 nW, which correspond to photon flux densities of 0 s^−1^µm^−2^, 3.1 × 10^6^ s^−1^µm^−2^, and 6.5 × 10^6^ s^−1^µm^−2^. These backgrounds bleached 0%, 60%, and 85% of photopigment in the 10 s prior to the stimulus flash.

All patterns were employed for measurements on both subjects, with the exception of the ‘multiple flashes’ pattern, which was measured in only one of the subjects.

### 4.5 Signal processing

The acquired spectral stacks were processed using a procedure described in detail elsewhere (*23*). In short, each acquired interferogram modulated by the off-axis carrier frequency was demodulated by 2D Fourier transform in *xy*, filtering in the Fourier space, shifting, and 2D inverse Fourier transformation. Next, a one-dimensional short-time Fourier transformation in the spectral dimension was used to estimate fringe chirp due to system vibrations and/or group velocity dispersion mismatch. This chirp was then corrected by adding phase to the spectral stack accordingly. Finally, the corrected spectral stacks were Fourier transformed into OCT volumes.

The acquired volumes underwent a flattening process to correct for the tilt caused by the off-axis approach. This flattening was performed by fitting a plane to the surface of the retina and shifting pixels accordingly. The volumes were then segmented axially and the photoreceptor inner segment - outer segment junction (ISOS) and cone outer segment tip (COST) layers were identified and aerially projected to produce *en face* images. These projections were registered to one another in the two lateral dimensions using cross-correlation to produce a trace of lateral retinal motion during image acquisition. The *en face* registered images are then averaged and cones were automatically identified and individually segmented in three-dimensions. The cone row-to-row spacing at the 2° imaged location was approximately 4.5 µm, and center-to-center spacing thus 5.2 µm. Using the axial and lateral segmentation coordinates for each cone and the lateral eye movement trace, a volumetric region of interest (“subvolume”) was extracted for each cone from each OCT volumetric image in the series. For each cone, ORG processing was performed on this series of cone subvolumes.

From each subvolume, nine A-scans near the center of the cone were analyzed, spanning 2.7 µm. Each A-scan contained signal from the ISOS and COST surfaces. The phase difference between them at any given time (ΔΦ(*t*)) was calculated by computing the product of the complex COST pixel (*A*) and conjugate of the complex ISOS pixel (*B*) in each of the nine A-scans in the subvolume, computing the vector sum of these products, and finally computing the angle of the vector sum:

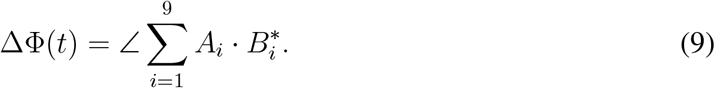

The phase difference was then converted to optical path length difference by

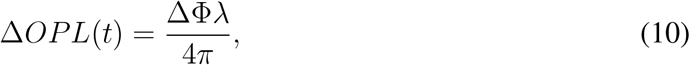

where *λ* is the wavelength of the imaging beam.

### 4.6 A predictive model for light-evoked OS deformations

In trials in which the stimulus was a single flash, the resulting time series of phase differences exhibited a fast contraction immediately after the flash onset, followed by a slower elongation, and an even slower recovery to baseline length.

The early contractile response of the OS has been attributed to the early receptor potential– a change in charge distribution around opsin proteins following isomerization of their retinal chromophores by light (*14*). This electrical shift is hypothesized to cause repulsion among the membrane-bound opsins, which in turn flattens the discs and contracts the OS (*20*). The slower and larger magnitude elongation that follows has been shown to be suppressed in a transducin-knockout mouse model (*27*), and has been hypothetically attributed by several groups to swelling of the OS associated with increasing intracellular osmolarity. Quantitative characteristics of these responses, such as maximum and minimum excursion and initial elongation velocity have been shown to depend on bleaching level by us and others (*10, 12–16*).

To provide a quantitative description of the initial contraction, subsequent elongation, and recovery toward baseline, a four-parameter exponential model was employed (Eq. 1):

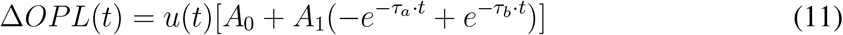

The four free parameters in this model are amplitudes *A*_0_, *A*_1_ and time constants *τ*_*a*_, *τ*_*b*_. Time is represented by *t*, with flash onset at *t* = 0, and Δ*OPL*(*t*) represents the change from baseline in OS optical path length at time *t. u*(*t*) is the Heaviside step function, defined as:

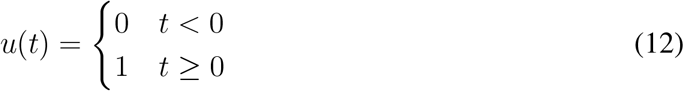

The data acquired for various stimulus levels were fitted to equation 1 in MATLAB (MathWorks, Nattick, MA) with a trust-region fitting algorithm (*28*). The parameters *τ*_*a*_ and *τ*_*b*_ were constrained to be non-zero positive values, to prevent divergence of Δ*OPL* when *t* → ∞. The fast contraction of the OS, because it occurs over only a few experimental data points, contributes a relatively small (1% to 3%) component the summed square error (SSE) during curve fitting. To ensure that this part of the response was accurately captured, data points in this region were weighted by a factor of 10, such that squared error in this region was multiplied by 10 prior to summation of SSE.

Optimization of ORG methods for clinical applications requires us to know how the duration of measurement affects estimation of model parameters. Longer measurements presumably provide more repeatable parameter estimation, but at the expense of data processing and storage costs, especially because the data rate of the system is > 10 GB/s. To determine the effect of the duration of measurement on subsequent fitting, full 3 s measurements from subject 1 were fit with the model, and then truncated subsets of the data were fit with the same model. The effect of varying the data length on model parameters was expressed as a percentage error (*PE*) in fitting parameters (*p*) as a function of the data duration, with the full dataset (3 s observation window) used as the reference (*p*_*r*_):

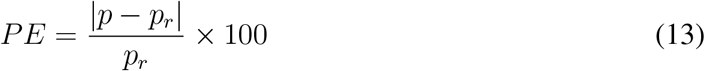

For the proposed model to have predictive utility, i.e., to help predict the response of cones to arbitrary stimuli, the parameters of the model must be described as functions of stimulus dose. To achieve this, the relationships between model parameter estimates and dose were fitted as well, using log-linear and Michaelis-Menten equations.

To quantify responses to two flashes (referred to as *α* and *β*) separated by an inter-stimulus interval of *t*_isi_, responses were fit with a time-shifted sum of two single-flash models, described in Eq. 14. Quantifying responses to multiple flashes permits assessment of the additivity of single flash responses.

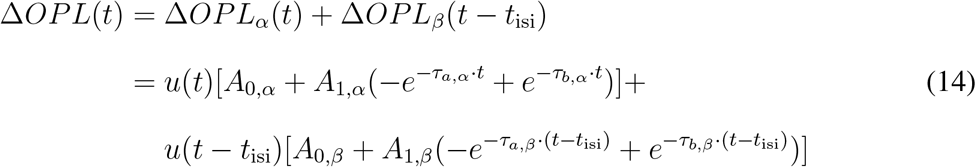

To quantify responses to flashes delivered after subjects had adapted to a dim background for 10 s, we modeled the response as a sum of the single flash response (Eq. 1) and a linear ramp *m* · *t*, where the slope of the ramp *m* is the rate of photopigment bleaching due to the background. The acquired data were fit to the following equation:

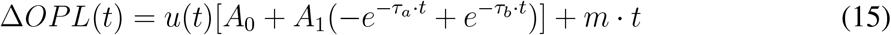

## Acknowledgements

The authors acknowledge the assistance of clinical coordinator Susan M. Garcia and helpful discussions with Jack Werner PhD, Ewelina Pijewska PhD, Maciej Bartuzel PhD, Ratheesh Meleppat PhD, and Donald T. Miller PhD.

## Funding

This work was supported by National Institutes of Health (R01-EY-034340, R01-EY-033532, R01-EY-031098, R01-EY-026556, P30-EY-012576), Research Council of Finland (356826).

## Author Contributions

RSJ conceived the research. DV and RJZ designed the apparatus. DV constructed the apparatus. DV and RSJ developed the instrumentation software. RJZ, RSJ, and DV designed the signal processing pipeline. DV and KVV conducted the experiments. DV conducted the signal processing. All authors designed and conducted the data analysis. All authors wrote the manuscript.

## Competing Interests

The authors declare that they have no competing financial interests.

## Data and materials availability

Additional data and materials will be made available upon reasonable request.

## Notes

### Competing Interest Statement

The authors have declared no competing interest.

