## Supplementary materials for "Insight into human photoreceptor function: modeling optoretinographic responses to diverse stimuli"

Denise Valente et al.

**This PDF file includes:**

1. Tables S1-S3.
2. Figures S1-S3.

### 1 Tables

Table S1: Goodness of fit ($R^2$) and residual RMS error (RMSE) for all measurements.

| | Subject 1 | | Subject 2 | |
|---|---|---|---|---|
| Bleaching % | $R^2$ | RMSE (nm) | $R^2$ | RMSE (nm) |
| 1 | 0.9635 | 2.7165 | 0.9587 | 3.3486 |
| 2 | 0.9818 | 2.7511 | 0.9809 | 3.2190 |
| 4 | 0.9934 | 2.9354 | 0.9926 | 3.2291 |
| 8 | 0.9889 | 6.4219 | 0.9898 | 6.0038 |
| 16 | 0.9966 | 5.7911 | 0.9937 | 6.8220 |
| 32 | 0.9979 | 9.6678 | 0.9953 | 11.0914 |
| 64 | 0.9967 | 16.8443 | 0.9917 | 21.7070 |

Table S2: Residual RMS fitting error (RMSE) for various models. Line colors refer to plots shown in main paper, Fig. 10.

| Model equation[a] | Line color | RMS error | RMS error $t \geq 0.5\,\mathrm{s}$ |
|---|---|---|---|
| $A_0 + A_1\left[-e^{-\tau_a \cdot t} + e^{-\tau_b \cdot t}\right]$ | magenta | $3.5\,\mathrm{nm}$ | $3.0\,\mathrm{nm}$ |
| $A_1\left[-e^{-\tau_a \cdot (t-t_0)} + e^{-\tau_b \cdot (t-t_0)}\right]$ | green | $3.5\,\mathrm{nm}$ | $3.0\,\mathrm{nm}$ |
| $A_1\left[-e^{-\tau_a \cdot t} + e^{-\tau_b \cdot t}\right]$ | blue | $6.7\,\mathrm{nm}$ | $4.7\,\mathrm{nm}$ |
| $A_1\left[1 - e^{-\tau_a \cdot t}\right]$ | red | $8.2\,\mathrm{nm}$ | $4.0\,\mathrm{nm}$ |
| $A_0\left[1 - e^{-\tau_a \cdot t}\right] + A_1\left[1 - e^{-\tau_b \cdot t}\right]$ | cyan | $8.2\,\mathrm{nm}$ | $4.0\,\mathrm{nm}$ |

[a] For simplicity, multiplication by the Heaviside function $u(t)$ is omitted.

Table S3: Photopigment bleaching levels

| Bleaching percentage | Power ($\mu W$) | Pulse width (ms) | Photons/s | Photon count | Photon density (photons/$\mu m^2$) |
|---|---|---|---|---|---|
| 1 | 1.25 | 10 | 3.49E+12 | 3.49E+10 | 3.43E+5 |
| 2 | 2.5 | 10 | 6.98E+12 | 6.98E+10 | 6.86E+5 |
| 4 | 5.05 | 10 | 1.41E+13 | 1.41E+11 | 1.39E+06 |
| 8 | 10.3 | 10 | 2.88E+13 | 2.88E+11 | 2.83E+06 |
| 16 | 21.5 | 10 | 6.01E+13 | 6.01E+11 | 5.90E+06 |
| 32 | 47.8 | 10 | 1.34E+14 | 1.34E+12 | 1.31E+07 |
| 64 | 100.2 | 12.6[c] | 2.80E+14 | 3.53E+12 | 3.47E+07 |

### 2 Figures

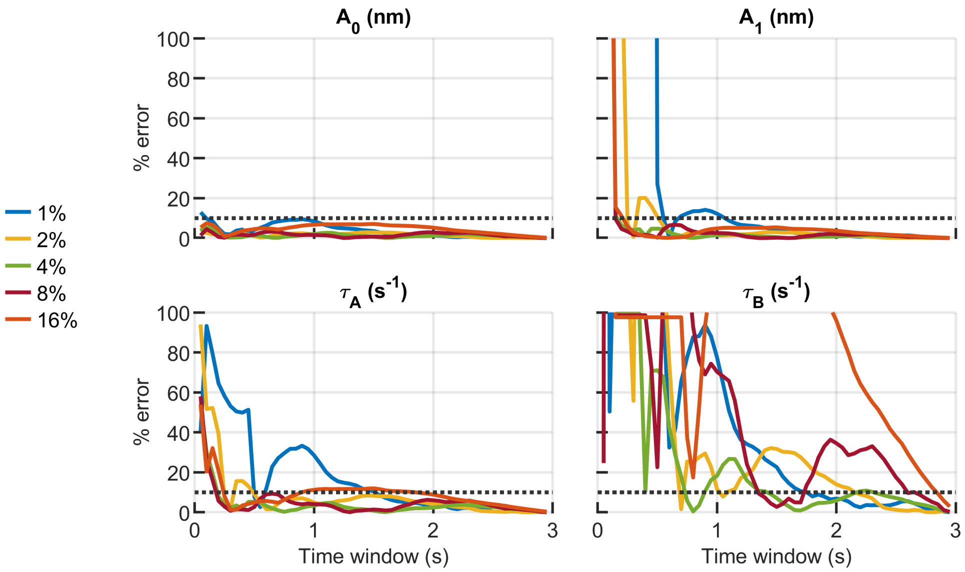


Figure S1: **Parameter estimation and measurement duration.** Over a range of bleaching percentages $1\,\%$ to $16\,\%$, the model was fit to subsets of measured data, and resulting estimates of each parameter were compared to the estimate derived from the full, uncropped data set and expressed as a percent error. The biggest variations occurs for $\tau_B$, the parameter associated with the slow return of OS length to baseline. These findings imply that measurement durations longer than $3\,\mathrm{s}$ may be required to assess this recovery step. A threshold of $10\,\%$ was selected as an acceptable percentage error (black dashed line).

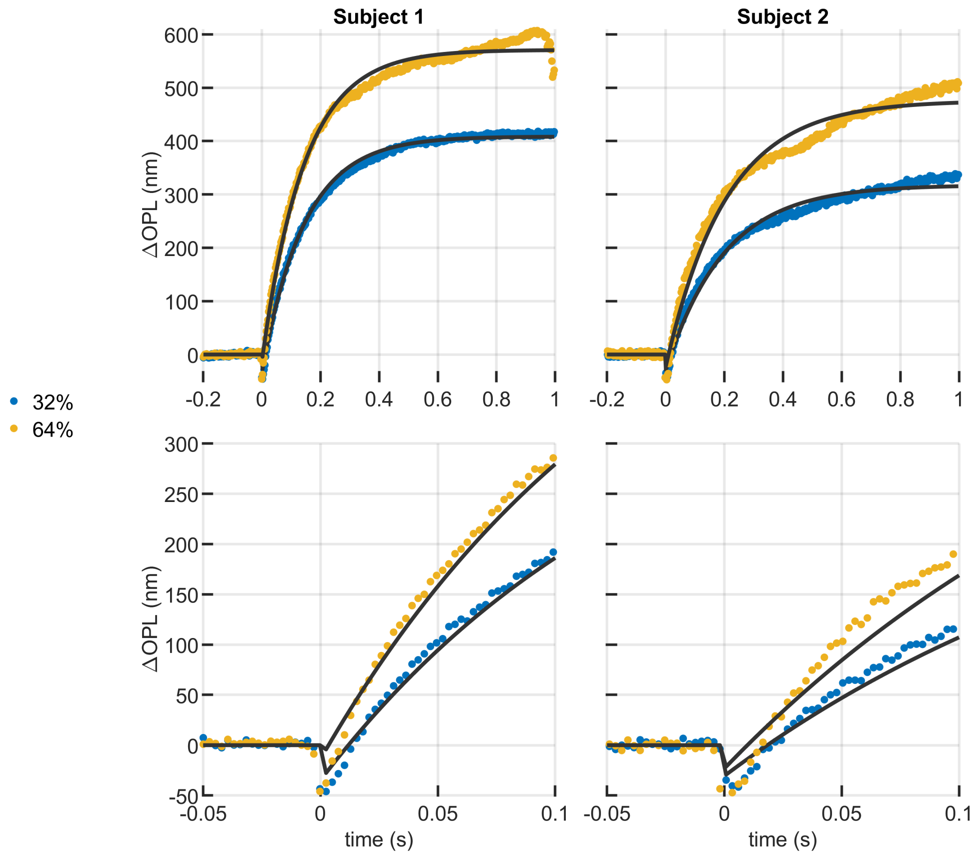


Figure S2: **Curve fitting of ORG responses at higher bleaching levels.** (top) It can be seen that at high bleaching levels the model starts to diverge from the experimental data at both the swelling (top) and contraction (bottom) stages. This divergence could indicate that either another exponential component with smaller amplitude might be present or that nonlinear biomechanical factors influence the response.

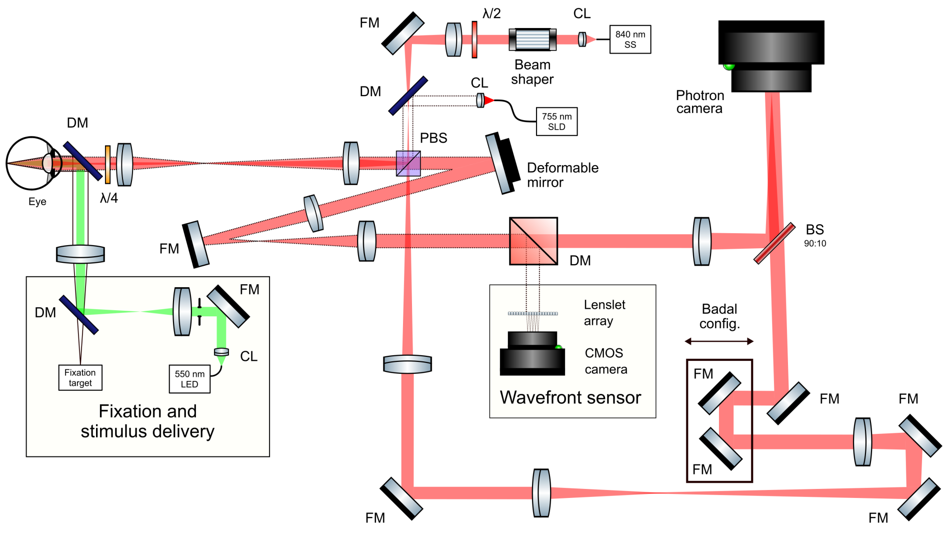


Figure S3: **Schematic of the AO-FF-SS-OCT setup.** The tunable light source for imaging is represented in solid red, while the red dashed beam represents the beacon for AO operation. The beam represented in green is the LED to deliver the stimulus. CL: collimation lens, $\lambda/2$: half-wave plate, $\lambda/4$: quarter-wave plate, FM: flat mirror, DM: dichroic mirror, BS: beam splitter, PBS: polarizing beam splitter. (Schematic not drawn to scale.)
